# Local adaptation and maladaptation during the worldwide range expansion of a selffertilizing plant

**DOI:** 10.1101/308619

**Authors:** A. Cornille, A. Salcedo, H. Huang, D. Kryvokhyzha, K. Holm, X-J Ge, J.R. Stinchcombe, S. Glémin, S.I. Wright, M. Lascoux

## Abstract

Species having experienced rapid range expansion represent unique opportunities to evaluate the dynamics of adaptation during colonization of new environments. We investigated the consequences of range expansion on local adaptation of a successful worldwide colonizer, the shepherd’s purse *Capsella bursa-pastoris*. This species is an annual weed that originated recently in Eurasia and has now broadly colonized both temperate and subtropical areas. We assessed the performance, genetic diversity, and phenology of field-collected accessions belonging to three distinct genetic clusters of decreasing age (Middle East, Europe and Asia) in three common gardens in Europe, Asia and North America. To understand the genetic basis of local adaptation in this species, we also tested for correlation between SNP allele frequencies and environmental factors in Europe and Asia. Overall, we showed that patterns of local adaptation depended on population history: some older populations were weakly adapted to local conditions while those closer to the front of the colonization wave, far from the origin of the species, were maladapted whatever the common gardens. Altogether, our results have important consequences for the understanding of the evolution and adaptation of self-fertilizing plant during range expansion.

## INTRODUCTION

Successful range expansion inherently depends on the ability of species to rapidly adapt to new local environmental conditions. Understanding the evolutionary processes underlying a species distribution is, therefore, closely related to understanding local adaptation. While we are beginning to have a fairly good grasp of the forces involved in local adaptation for populations at equilibrium, the extent to which more marginal and recent populations characterized by non-equilibrium demographic history are expected to be locally adapted remains unclear. There are many reasons to believe that marginal populations may not be locally adapted and could even be maladapted. First, theory predicts that gene flow from central to marginal populations would lead to maladaptation in the latter (Haldane 1957). Second, theory also predicts that range expansion can enable both favorable and deleterious mutations to establish at expanding range margins and reach high frequencies, a phenomenon called surfing, which in the case of deleterious alleles creates a so-called ‘expansion load’ (Peischl *et al.* 2013, 2015, 2018; Peischl & Excoffier 2015). If expansion loads occur frequently, maladapted genotypes and phenotypes can, therefore, spread over the wave edge for thousands of generations (Peischl *et al.* 2015). This is especially true in self-fertilizing (selfing) species. The ability to self confers an advantage over outcrossing species since a single individual can colonize a new environment (Baker *et al.* 1965), a phenomenon referred to as reproductive assurance, but it also implies that colonization may proceed through a series of severe bottlenecks, leading to an accelerated loss of genetic variation and decline in fitness. Hence, the apparent striking success of some selfers in colonizing the world may actually be short-lived and, over longer timescales, could accelerate their demise. Park and co-authors recently argued that, in general, selfing species, even those that have successfully colonized large territories, actually are exhibiting diminished niche breadth over time (Park *et al.* 2017). Understanding the pace at which local adaptation is established in rapid colonizers is, therefore, a core problem in evolutionary biology with both fundamental and applied consequences; this information is indeed crucial to forecast how species are likely to respond evolutionarily to rapid global environmental changes (Bellard *et al.* 2012).

Classical studies investigating the amount of local adaptation are based on reciprocal transplant experiments measuring the fitness and phenotypes of populations in their own habitat and when transplanted to other habitats (Turesson 1922; Swanson 1949; Antonovics & Bradshaw 1970; Gómez *et al.* 2009; Blanquart *et al.* 2013). Or, when such reciprocal transplantation is not possible, local adaptation can be evaluated on common garden experiments, where different populations are tested under the same set of environmental conditions in one or more locations (Savolainen *et al.* 2013). Reciprocal transplant experiments are among the most powerful tools to investigate the genetic basis of local adaptation. However, reciprocal transplant experiments also have significant limitations because they are generally restricted to a small number of populations and usually do not incorporate the demographic histories of different populations. Yet, taking into account population demographic histories is crucial when analyzing the evolutionary causes of fitness differences among populations (Tiffin & Ross-Ibarra 2014). While it is tempting to ascribe differences in individual fitness and phenotype variation to adaptive evolution, evidence of selection should be evaluated while accounting for neutral evolutionary processes since random events can also affect individual fitness and quantitative traits. With the advent of high-throughput sequencing and the development of large-scale inference methods over the past decade, it is possible to reconstruct fairly well detailed demographic histories for a species (Knowles & Alvarado-Serrano 2010; Sousa *et al.* 2014; Schraiber & Akey 2015; Maisano Delser *et al.* 2016). This can provide neutral expectations that can then be incorporated in the study of local adaptation (Tiffin & Ross-Ibarra 2014). Such neutral expectations can be efficiently integrated by a series of common garden experiments located to reflect the species’ demographic history. In particular, in the case of a species having experienced a range expansion, characterizing individual fitness and phenotypes of the species with multiple common gardens across the colonization range is an elegant approach to investigate the impact of range expansion on the pattern of local adaptation.

Here, we investigated the level of local adaptation in a successful worldwide colonizer of allopolyploid origin (Douglas *et al*. 2015), the shepherd’s purse, *Capsella bursa-pastoris*, whose demographic history was recently inferred from genome-wide markers (Cornille *et al*. 2016). A recent population genomic study within the ancestral range of *C. bursa-pastoris* (Cornille *et al*. 2016) revealed three distinct genetic clusters: in Asia (ASI), in the Middle East and northern Africa (ME) and in Europe and the Russian Far East (EUR). Demographic inferences also showed that these three clusters are the result of a recent expansion that likely started from the Middle East (ME) and was followed by two consecutive colonization events, first towards Europe (EUR) and then, more recently, into Asia (ASI). Hence, considering that *C. bursa-pastoris* originated around 100,000 years ago (Douglas *et al*. 2015) and was affected by glacial cycles, its current extensive distribution should be the result of a recent and rapid range expansion. We also showed that the three clusters differed markedly in genetic load, with a much higher load in Asia than in Europe or in the Middle East (Kryvokhyzha *et al*. 2017), and that the global pattern of gene expression between the three main clusters primarily reflected the *C. bursa-pastoris* demographic history (Kryvokhyzha *et al*. 2016).

To assess the impacts of demographic history, in particular of rapid range expansion, on local adaptation, we established three common gardens of 257 accessions to investigate phenotypic and fitness variation of *C. bursa-pastoris*. The originality of our experimental design relies on the use of a comprehensive sample of populations that can be a good proxy of *C. bursa-pastoris* metapopulation dynamics in Eurasia, a critical criterion to optimize the power to detect local adaptation (Blanquart *et al*. 2013). These three common gardens were installed at two latitudinal extreme locations in Eurasia, one in East Asia and the other in Northern Europe, as well as in North America which is out of the native range of the species. We complemented our common garden experiments with environmental association scans to look for subtler evidence of adaptation by associating genotyping by sequencing (referred to as GBS hereafter) SNP frequencies (Cornille *et al*. 2016) to publicly available climate variables.

Using these complementary approaches, we tested for evidence of local adaptation or expansion load by addressing the following questions: What is the individual fitness across the three transplant sites? Does genetic origin (ASI, EUR, ME) contribute to fitness differences that could be linked to local adaptation or expansion load? Has adaptation to different environmental conditions yielded differences in putatively adaptive traits (flowering time and floral display)? Are SNPs associated with climatic variables? Does genetic origin impact the genetic architecture of adaptation? Our data supported a weak signal of local adaptation in the oldest *C. bursa-pastoris* populations (ME) and of maladaptation in the youngest one (ASI). Complementary environmental association analyses allowed for identifying possible climatic adaptation in the genome without advanced knowledge of the traits involved and detect signals of adaptation beyond the conditions incorporated in the common gardens. Altogether, this study allows us to understand how contrasting demography influences adaptation at the genetic and phenotypic levels.

## Materials and methods

### 1. Plant material

The plant material consisted of 257 accessions collected across 64 populations in Eurasia (Table S1 and Figure 1A) that represent different environments (photoperiod, temperature, precipitation, humidity etc.) (Kryvokhyzha *et al*. 2016). For each accession, seeds were collected from individual mother plants sampled in the field and were grown for one generation under controlled conditions in growth chambers to avoid maternal effects. Almost all accessions have been genotyped using GBS (Cornille *et al*. 2016), as well as phenotyped for flowering time under controlled conditions (Kryvokhyzha *et al*. 2016). These accessions are known to belong to three main genetic groups in Eurasia: Europe (EUR), Middle East (ME) and Asia (ASI) (Cornille *et al*. 2016). Russian accessions were previously assigned to the EUR cluster (Cornille *et al*. 2016). We therefore performed statistical analyses on the entire dataset, as well as on each genetic group considered separately (EUR, ASI and ME datasets hereafter). All accessions are listed in Figure 1A and Table S1 with their respective assignation to each genetic cluster.

**Figure 1.**
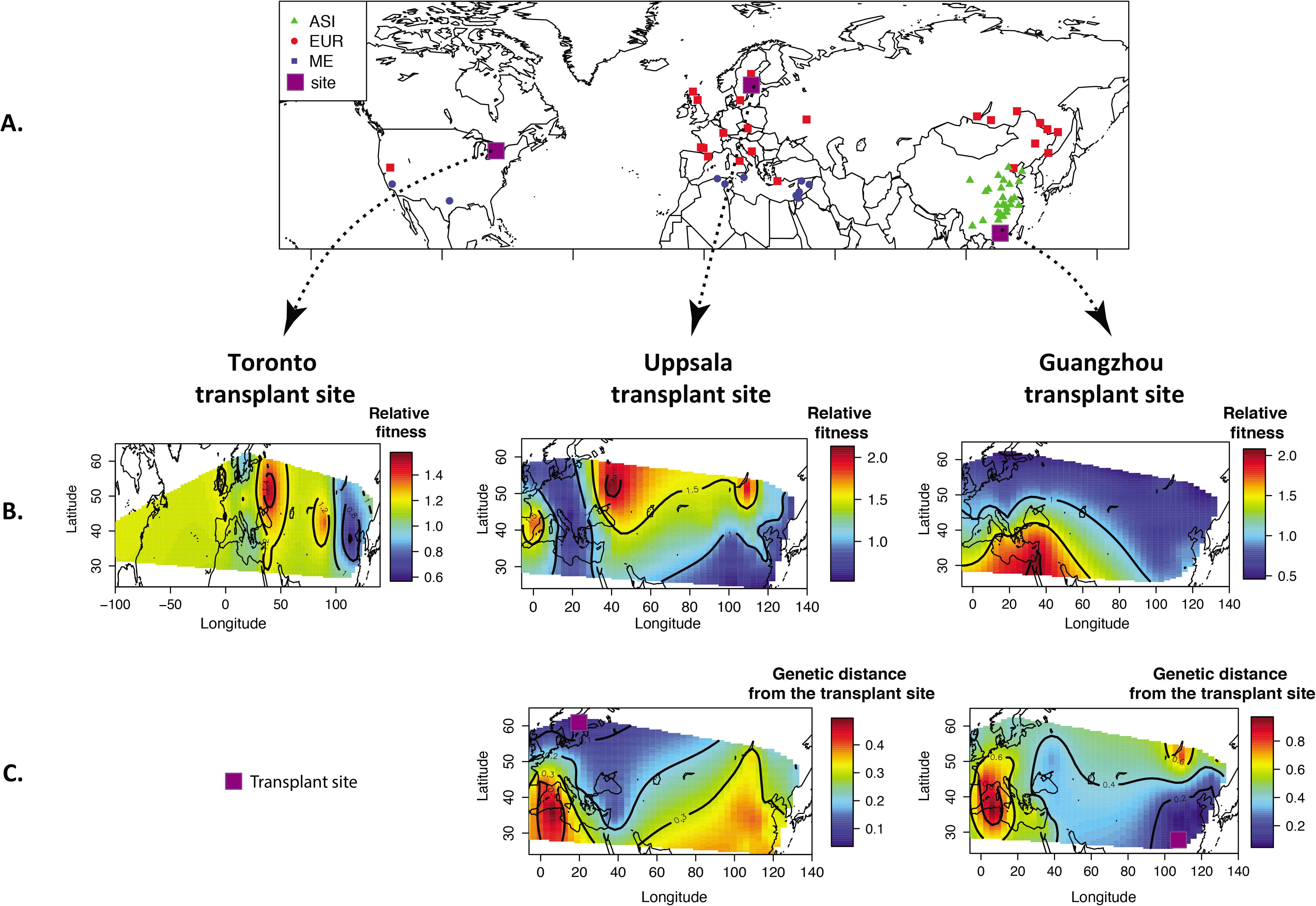
**A.** Geographical origins of the 64 sites of *Capsella bursa-pastoris* used in this study, with their respective genetic cluster assignation. Square in purple represented the three common gardens in Uppsala (Sweden), Toronto (Canada) and Guangzhou (China). **B.** Map of interpolated mean individual relative fitness for each common garden (upper part) and **C.** interpolated mean genetic distance per accession from the Guangzhou and the Uppsala common gardens (interpolation for the TOR dataset is not presented because no genetic information for this place was available, Cornille et al. 2016). ASI: Asian genetic group, EUR: European genetic group, ME: Middle Eastern genetic group.

### 2. Transplant experiment

#### 2.1. Design of the transplant experiment

Seeds of the accessions were transplanted in common gardens at three geographical sites: Uppsala (Sweden, 59°N), Toronto (Canada, 43°N) and Guangzhou (China, 23°N). The three sites were chosen to represent different climatic conditions (Figure S1): Uppsala and Guangzhou represent two contrasting climates within the natural range of *C. bursa-pastoris*, whereas Toronto represents a site outside of the native distribution of *C. bursa-pastoris*, with environmental conditions most closely resembling those of the Uppsala site (Figure S1). Depending on the seed germination success of the 257 accessions, the final number of accessions per transplant site was variable, but all three genetic clusters were well represented at each site (Table S1). To benefit from proper climatic conditions, the experiments were installed in early May 2014 in Uppsala, early June 2014 in Toronto and early November 2014 in Guangzhou. During the experiment, we also recorded mean temperature, mean humidity and day length in each transplant site.

The experiments were organized in six blocks, each including one individual per accession. The accessions were randomly distributed within each block. Each block included 231 and 257 accessions in Guangzhou and Uppsala, respectively, corresponding to a total of 1,386 and 1,542 plants, respectively. A reduced common garden was also installed in Toronto: the experiment comprised 160 accessions organized in five blocks representing a total of 800 plants. At the Toronto site, we carefully subsampled the accessions by maximizing genetic diversity. To that end, we first estimated genetic similarity using identity by state and only retained lines with less than 97% similarity.

In each site, about 20 seeds per accession were sterilized all the same day and germinated in petri dishes, with MS medium and Agar (see details of the protocol in (Kryvokhyzha *et al*. 2016). Petri dishes were then stratified for seven days at 4°C in the dark in order to promote germination. After this seven-day cold treatment, petri dishes were placed in a greenhouse to protect seeds from rainfall. The conditions within the greenhouse mimicked the outdoor conditions, with no additional light or heating. Germination was monitored daily for each replicate of each accession. Petri dishes were randomized and moved every day to avoid micro-environmental variation effects.

In Uppsala and Guangzhou, a few seedlings per accession were transplanted into individual pots and thinned to one per pot when they reached a four-leaf stage (16-20 days after cold treatment). One week after being thinned, pots were then placed outside in a common garden. The six (or five) blocks (1m x 3m, 9 x 30 arrays) were arranged at 2m spacing and aligned longitudinally. The bottom of the pots was pierced so that roots were able to grow outside of the pots easily. In Toronto, seedlings were transplanted directly into 13’ x 5’ beds containing 5cm of ProMix soil, without pots. The plants were watered twice the first week to ease acclimatizing to outdoor conditions, and at Toronto the plants were shaded for two days after planting.

#### 2.2. Phenological, morphological and fitness measurements

Plants were monitored daily. Over the course of each experiment, we monitored four phenological traits (Table 1): germination time, bolting time (i.e. differentiation of the bud from vegetative parts indicating the initiation of the reproductive period), flowering time (i.e. number of days from germination to the appearance of the first open flower) and reproductive period (i.e. number of days from the appearance of the first flower until the senescence of the plant). At flowering time, we also recorded two phenotypic traits: the rosette leaf number and the maximum diameter of the rosette (Table 1).

**Table 1.** Abbreviations used in the manuscript.

| Abbreviations | Full names |
| --- | --- |
| EUR | European genetic group detected using the 4,274 SNPs dataset |
| ME | Middle Eastern genetic group detected using the 4,274 SNPs dataset |
| ASI | Asian genetic group detected using the 4,274 SNPs dataset |
| UPP | Common garden built in Uppsala (Sweden) |
| GUANG | Common garden built in Guangzhou (China) |
| TOR | Common garden built in Toronto (Canada) |
| GT | Germination time |
| BT | Bolting time |
| FT | Flowering time |
| RP | Reproductive period |
| LEAVESNB | Rosette leaf number at flowering |
| ROSDIA | Maximum diameter of the rosette |
| W and w | Absolute and relative fitness, respectively |

To assess plant fitness, we recorded the height (in cm) of the highest inflorescence at the time of senescence of the plant, the number of basal inflorescences and the number of secondary inflorescences on the main stem. We then counted the number of fruits along a 10cm section in the middle of the main inflorescence. Unlike in *Arabidopsis* (Roux *et al*. 2004), the correlation of fruit size and the number of seeds has never been estimated in *C. bursa-pastoris*. We therefore also quantified the mean fruit width and the mean height of fruit incision as the depth of incision in mm between the two valves of the fruit on five fruits per plant. The mean number of seeds per fruit for a total of five fruits was then recorded for at least two out of the six (or five) replicates. For the Uppsala and Guangzhou datasets, the fitness for each plant was then calculated as the product of the number of fruits on a 10 cm section of the main inflorescence, the height of the main inflorescence divided by 10 cm, the number of primary branches, assumed to be approximately the same length and carry the same number of fruits as the main branch, and the mean number of seeds per fruit. For the Toronto dataset, the fitness for each plant was calculated as the product of the mean number of seeds per fruit, the number of fruits on the main inflorescence, and the number of primary branches.

#### 2.3. Spatial pattern of local adaptation

Before testing explicitly for a signal of local adaptation, we first described the spatial pattern of mean relative fitness per accession across Eurasia for each transplant site with the *kriging* and *surface* functions (*fields* R package) (Nychka *et al*. 2017).

We first formally tested for local adaptation by considering all available populations, which essentially amounts to consider all the sampled populations as a subset of a larger metapopulation. This approach allowed for estimating intrinsic site and population quality effects (Blanquart *et al.* 2013) and considered local adaptation as a characteristic of the whole metapopulation.

We first combined the three transplant site datasets (i.e. equivalent of three replicates of the experiment in different environments) to test for the difference between the fitness of populations in their own site (in sympatry) and the fitness of populations transplanted to other sites (in allopatry). To test this sympatric *vs*. allopatric contrast, we used a generalized linear mixed model with a negative binomial error distribution and log-link function (*glmer.nb* function in the *lme4* R package) (Bates *et al.* 2015). The model was the following:

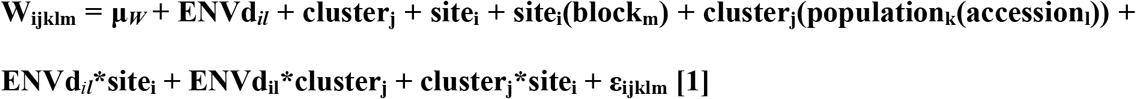

where *W_ijklm_* is the absolute fitness value of an individual in block *m* from accession *l* of population *k* in site *i, μ_W_* is the mean absolute fitness, *site_i_* is the common garden location (Uppsala, Guangzhou, Toronto), *ENVd_il_* is the environmental distance of accession *l* from site *i, block_m_* is the block effect within each site, *cluster_j_* is the genetic group (EUR, ASI, ME) and *ε_ijklm_* is the residual term. *Block* is random and nested within site, *population* is nested within cluster, and *accession* is nested within *population*, and they were added to the models as random-effect terms. To estimate *ENVd*, we calculated environmental distance of each accession to a given common garden, which was calculated based on 19 bioclimatic variables extracted from the Worldclim database (http://worldclim.org); these variables characterize the bioclimatic environment of the site of origin of each accession and of the three transplant sites (Text S1). The environmental distance was scaled and centered using the R function *scale* (see Supplementary Material for details, Text S1). We interpret *ENVd* as a measure of the strength of local adaptation (i.e. accessions from more environmentally divergent sites are predicted to have lower fitness under adaptation). The *site* term measures the quality or suitability of the transplant locations, *cluster* accounts for differences in fitness intrinsic to each genetic clusters, and the *ENVd_il_*cluster_j_* accounts for differences in local adaptation among the three genetic clusters. A full model (all main factors and two-way interactions) allowed us to test for local adaptation by quantifying the sympatric-allopatric contrast (Blanquart *et al.* 2013).

Because we detected a significant site*genetic cluster interaction, we then fit separate models to test the sympatric-allopatric contrast within each genetic cluster. We used a linear mixed model with a negative binomial error distribution and log-link function for each genetic cluster:

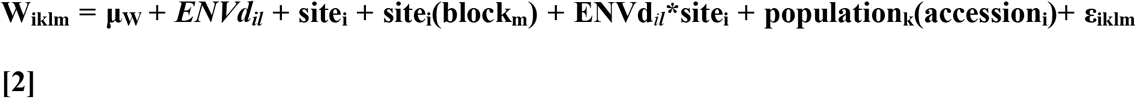

where *W_iklm_* is the absolute fitness of an individual plant from accession *l* of population *k* in *block m* of site *i, μ_W_* is the overall mean fitness, *site_i_* is the common garden (Uppsala, Guangzhou, Toronto), *ENVd_il_* is the environmental distance between accession *l* and site *i* (see supplementary material for details, Text S1), *block* is the block effect within each site and *ε_iklm_* is the residual term. *Block* is random and nested within *site. Accession* is nested within *population* and was added to the models as a random-effect term.

Finally, we investigated the fitness stability, or responsiveness, of each accession to environmental changes by using the Finlay-Wilkinson regression (FWR) (Finlay & Wilkinson 1963). With this method, the fitness of individual accessions over the three sites was regressed over the environment of the different sites, the quality of the environment of each site being simply characterized by the overall mean fitness of all accessions. We used the R package *FW* (Lian & de los Campos 2016) that implements a Bayesian FWR, with default parameters and the Gibbs sampling method. The FWR package allows the incorporation of the genetic relationship among accessions. We therefore used the genotypic relationship matrix based on the 4,274 SNPs obtained by Cornille *et al*. (2016). For each accession, FWR estimates a genotypic main effect and a slope (*b_i_*). The slope measures the linear response of an accession to the environment, relative to all other accessions in the dataset. A slope of one denotes an accession exhibiting the population average response to the environment, while a slope of zero denotes an accession that does not respond to the environment. A slope greater than one denotes an accession whose response to the environment is stronger than the average accession in the population (Kusmec *et al*. 2017). Conversely, a slope lower than one denotes an accession whose response to the environment is weaker than the average accession in the population. The data was analyzed using the entire dataset and for each genetic cluster (EUR, ME, ASI).

#### 2.4. Relationship between fitness and overall genetic diversity

We next investigated whether individuals from the front edge, which tend to have lower diversity and a higher number of deleterious mutations (Kryvokhyzha *et al*. 2017), also had lower fitness than those from more central populations, i.e. higher genetic load.

We investigated the relationship between fitness and genetic distance to the site where the fitness was assessed, which has seldom been explored (but see Montalvo & Ellstrand 2001; Hancock *et al*. 2011). We applied a principal component analysis (PCA) on the GBS dataset using the *dudi.pca* function from the *adegenet* R package (Jombart & Ahmed 2011) assuming two dimensions that represent each more than 5 % of the variance (first component: 19.8%; and second component: 8.3%). We then calculated absolute differences between each accession and the transplant site along the first two axes generated by a PCA as a proxy of genetic distances (*GENDist*). Since we did not have SNP characterization of accessions belonging to the exact location of each transplant site, we therefore used the genotype of the closest accession to the transplant site to compute the absolute differences. As the Toronto dataset was too far from the closest genotyped accessions (i.e. in the southern USA), *GENDist* was not analyzed for the Toronto dataset in model [3] below.

First, we tested whether the fitness of each accession decreased with increasing genetic distance (excluding the Toronto dataset), by taking into account the environmental distance as a covariate accounting for local adaptation. For each transplant habitat, we used a negative binomial error distribution and log-link function taking genetic distance (*GENDist*) as fixed factor and blocks (i.e. six blocks) and population of origin as random effects:

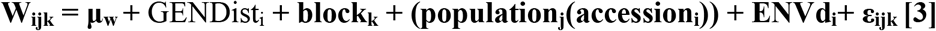

where *W_ijk_* is the absolute fitness of accession *i* from population *j* in block *k*.

We further explored signals of maladaptation by testing whether populations showing the lowest diversity, i.e. the ASI genetic cluster at the front edge, also showed the lowest fitness. To that end, we tested whether the fitness of populations belonging to each *C. bursa-pastoris* genetic cluster decreased with decreasing genetic diversity per population. For each common garden we ran a general linear model with a negative binomial distribution taking genetic diversity [π (Nei 1978)] per population, genetic cluster, and *ENVd* as fixed factors:

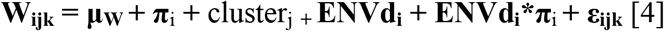

In this model, *W_ijk_* is the average absolute fitness of population *i* from cluster*j* in block *k*.

#### 2.5. Major phenotypic and adaptive differences among *C*. *bursa*-pastoris genetic groups

We assessed the strength and form of selection on different phenotypic traits and which ones could contribute to adaptive difference among genetic groups.

We first analyzed phenotypic differences between the three genetic groups and common gardens. We used a negative binomial error distribution and log-link function (*glmer.nb* function in the *lme4* R package) (Bates *et al.* 2015):

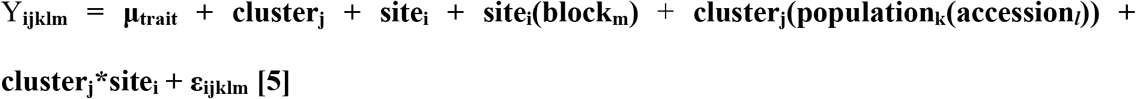

In this model, *Y* is one of the five phenological and morphological traits (i.e. BT, RP, FT, LEAVESNB, ROSDIA), *Y_ijklm_* is the phenotypic value of accession *l* belonging to population *k* and genetic cluster *j* grown in the block *m* at the site *i, μ* is the overall mean, *site* is the common garden (Uppsala, Guangzhou, Toronto), *block* is the block effect within each site, *cluster* is the genetic group (EUR, ASI, ME) and *ε* is the residual term. The same analysis, without the site effect, was also carried out for each common garden (model [5’]).

We next estimated selection on the phenotypic traits at each site, using selection differentials and gradients at each site (Lande & Arnold 1983; Arnold & Wade 1984). We used accession or genotypic means as our estimates of phenotypes and relative fitness, and as such selection differentials are the genetic covariance between traits and fitness (Robertson 1966), and gradients are genotypic selection gradients (Rausher 1992). Relative fitness for each accession at each site was estimated as the mean fitness value of the accession in the given site divided by the overall mean fitness value within the site (Lande & Arnold 1983; Arnold & Wade 1984; Stinchcombe *et al*. 2008). All traits were standardized by their within-site standard deviation to allow for comparison with other traits and sites. Although data deviated from a normal distribution (Shapiro-Wilk test, Table S2), these deviations were relatively minor as evidenced by the high Shapiro-Wilk statistic and visual examination of normal probability plots. Relative fitness was squared-root transformed to fit the assumption of normality.

Then, we estimated the selection gradients with the following models:

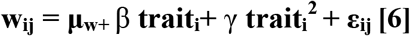

In these models, the partial linear regression coefficient (*β*) measures the strength and trend of directional selection and the quadratic regression coefficient (*γ*) measures nonlinear selection, which is suggestive of either stabilizing or disruptive selection. Following standard practices (Lande & Arnold 1983; Arnold & Wade 1984; Stinchcombe *et al*. 2008), the nonlinear selection coefficients presented here were doubled quadratic regression coefficients.

Finally, we tested differences in selection differential and linear (β) and quadratic (γ) coefficients among common gardens and among genetic clusters using analyses of covariance (ANCOVA). We modified our regression model above (model 6) by including main effects of site and genetic cluster, along with interactions between these main effects and the traits [model 7]. We interpreted a significant genetic cluster * trait, or site* trait interaction for relative fitness as evidence that selection on that trait differed by genetic cluster or site.

#### 3. Evidence of local adaptation through environmental associations using GBS

We performed environmental association analyses using the GBS data from 261 accessions (see Cornille et al 2016 for sequencing methods). To further reduce genotyping error rate below the reported 13% in Cornille *et al*. (2016), we further trimmed the GBS dataset (Text S2). After this filtering, we used a trimmed dataset of 3,801 SNPs with 8.6% error (Text S2) and publicly available climate data to test for environmental associations. As only 10 populations belonged to the ME cluster, and Middle Eastern and European populations showed evidence of admixture and shared ancestry (Cornille *et al.*, 2016), we grouped them with the populations from Western Europe to increase our power to detect SNP-environment associations throughout Europe. We tested environmental associations on the unphased GBS SNPs but removed all singletons. We also removed SNPs that were in >90% linkage disequilibrium (LD) within 10 SNP windows identified with the PLINK software (Purcell *et al*. 2007) to reduce multiple testing, and avoid detecting associations from the same locus through multiple tightly linked SNPs. After trimming and removing singletons, 1,567 and 1,513 SNPs remained in Asia and Europe, respectively. We compared our SNP data to a reduced set of publicly available climate data at 30 arc seconds (~1 km^2^) from the WORLDCLIM database (http://www.worldclim.org/). To reduce redundancy among the available environmental variables, we retained only variables correlated at *r*<0.8. We interpolated climate data for the GPS coordinates of each population using ArcGIS.

We first tested whether populations from more similar environments were more genetically similar by comparing genetic distance to climate distance. We summarized distance in climate by computing Gower’s distance among all populations using the 18 retained climate variables. We then tested for correlations between climate distance and *F_ST_* using a Mantel test with 1,000 permutations in the *vegan R* package (Oksanen *et al*. 2008). Next, we tested for SNP-environment associations with the individual-based Latent Factor Mixed Model (LFMM) approach (Frichot *et al*. 2013). The LFMM approach is similar to the mixed model regression approach often used in Genome Wide Association Studies (GWAS) and in landscape genomics as it also uses a regression to test how well environmental variables predict variance in genotypes while controlling for population structure at each SNP. However, LFMM shows higher power than traditional mixed model approaches (Frichot *et al.* 2013). We ran LFMM with the trimmed SNP dataset and the reduced set of environmental variables separately in both Europe and Asia. As recommended by Frichot *et al*. 2013, we used the number of significant eigenvalues computed through Tracy-Widom statistics in Eigenstrat as the dimensions of the latent factor matrix. We retained SNPs with the top 1% strongest associations. To investigate whether having closely related individuals within populations could lead to spurious associations, we also repeated the LFMM analysis using one randomly chosen individual per population. We then summarized SNPs with the 1% strongest associations from both approaches through their major PCA axes (eigenvalue greater than the mean eigenvalue), and compared how much variance in each of the environmental variables they explained. We found PCA axes summarizing significant SNPs from the full dataset explained more variance in their associated environmental variables than those summarizing SNPs identified from one individual per population in both Europe (All individuals mean *r*^2^=0.50; One individual per population mean *r*^2^=0.38, *P*=ns) and Asia (All individuals mean *r*^2^=0.61; One individual per population mean *r*^2^=0.49, *P*=0.051). We then carried out all subsequent analysis using SNPs identified from the entire sample.

Finally, we explored characteristics of the SNPs with significant associations. We tested whether we saw more SNPs and genes with significant associations in both Asia and Europe than expected by chance by repeating LFMM with 500 permutations of the environmental association data. We also compared the Minor Allele Frequency (MAF) of putatively adaptive SNPs to the MAF of significant SNPs from permutations to look for enrichment in allele frequencies within putatively adaptive SNPs. We then compared the z-scores of putatively adaptive SNPs, which reflect their effect size, between Europe and Asia, as well their LD. To evaluate whether LD for putatively adaptive SNPs in each region differed from that expected by chance, we compared mean pairwise *r^2^* among putatively adaptive to the distribution of mean pairwise *r^2^* from 1,000 random SNP draws of the same size as the test set from the corresponding trimmed GBS dataset. We computed LD both only among SNPs on the same scaffold, and among all SNPs. We next inquired about the origin of these SNPs by comparing the proportion of putatively adaptive SNPs in each region in each of the parental genomes through a *χ^2^* test.

## RESULTS

### 1. Spatial pattern of adaptation

The maps of interpolated mean fitness by population (Fig. 1B) for the three common gardens showed that accessions performed differently between common gardens. The maps indicated a better fit of South European and Russian accessions (i.e. the EUR genetic group) at the Uppsala common garden, a better fit of Middle Eastern accessions (i.e. the ME genetic group) at the Guangzhou common garden, and a better fit of the European and Middle Eastern accessions (the EUR and ME genetic groups) at the Toronto common garden. In all three common gardens, the Asian accessions (ASI genetic cluster) performed poorly compared to the accessions from the two other genetic clusters. We formally tested these observations hereafter.

### 2. Testing for local adaptation

The generalized linear mixed-effects model for the whole dataset (model [1]) showed a strong site*genetic cluster interaction (Table 2), indicating that the three genetic clusters performed differently in the three common gardens. However, we did not find any significant *ENVd* effect, indicating a weak effect of local adaptation at the population level when controlling for population structure. We therefore investigated patterns of adaptation testing within-genetic cluster fitness variation across sites (model [2], Table 2 and Figure 2). There was also a strong site effect but both *ENVd* and *ENVd*\*site effects were non-significant within each of the three clusters (Table 2). The ASI accessions showed consistently low fitness over the three sites, whereas the EUR accessions showed the greatest fitness at the Uppsala site and the ME accessions had the highest fitness at the Guangzhou site (Figure 2). Overall, the ASI accessions performed poorly at all sites, in Northern Europe the mean fitness of the EUR genetic cluster was greater than that of the ME, and the opposite was true at the Guangzhou site. At the Toronto site, which was outside of the original range of *C. bursa-pastoris*, the ME and EUR genetic clusters had similar mean fitness. These results suggested different signals of adaptation depending on the genetic clusters: general maladaptation for the ASI genetic group, and a signal resembling local adaptation, but non-significant while controlling population structure, for the ME and EUR genetic groups; ME had a higher fitness than EUR at the Guangzhou site, where environmental conditions are closer to the Middle East than in Northern Europe, and the reverse was observed at the Uppsala site.

**Figure 2.**
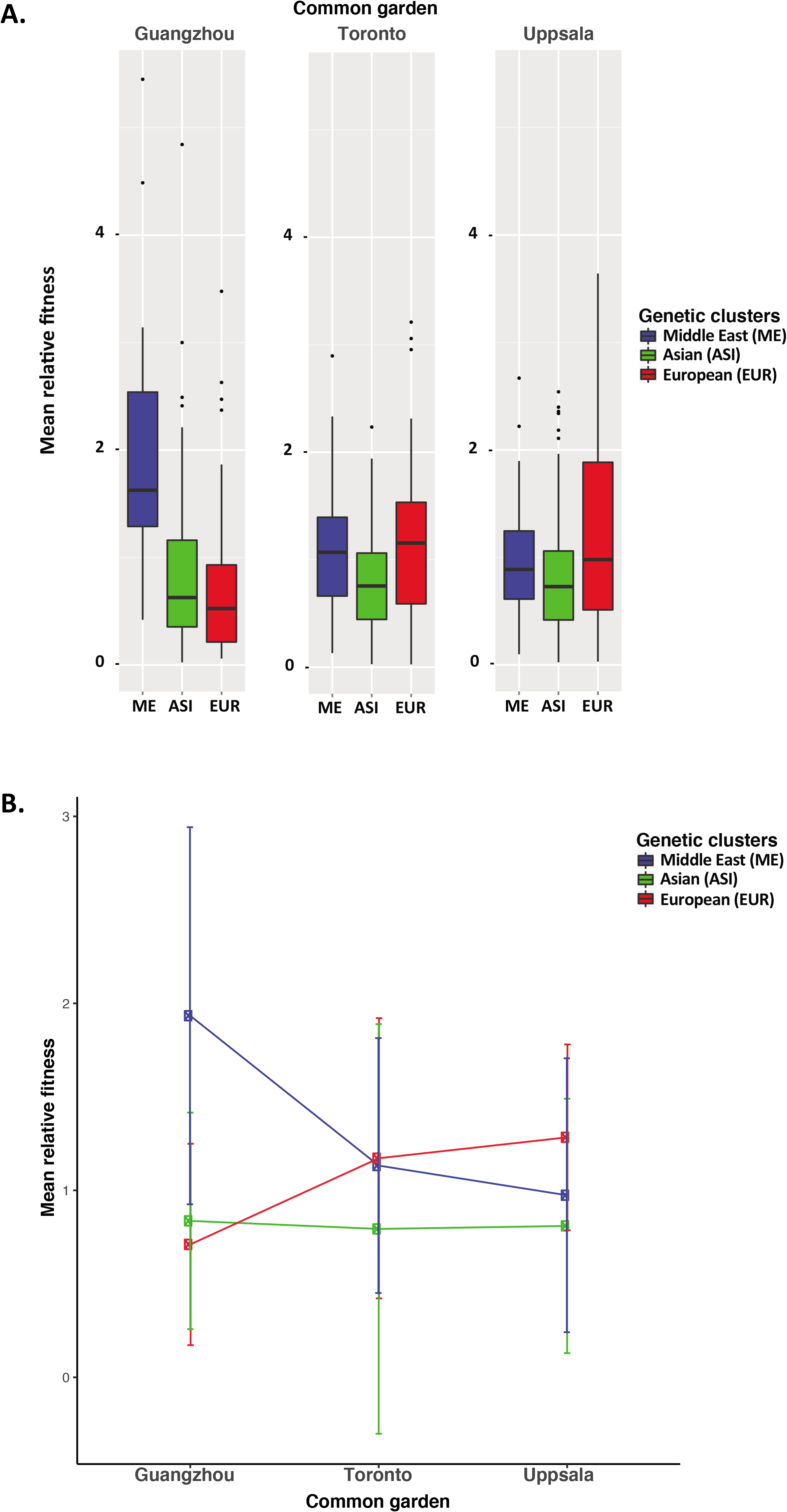
Performances of genetic groups in each common garden for *Capsella bursa-pastoris*. A) Within-genetic group performance (mean relative fitness per accession) at each common garden; B) Genetic group (EUR, ASI, ME) by site (Uppsala, Toronto, Guangzhou) interaction plot.

**Table 2.** Effects of the common garden site (site), the environmental distance (ENVd), the genetic cluster (EUR, ME or ASI) on the individual fitness (W) of the transplanted adult individuals of *Capsella bursa-pastoris* (Models [1] and [2]).

|  | W_WHOLE‡ |  |  | W_ASI‡ |  |  | W_EUR‡ |  |  | W_ME‡ |  |  |
| --- | --- | --- | --- | --- | --- | --- | --- | --- | --- | --- | --- | --- |
| Factors | Df | Estimate | $\chi^2$ | Df | Estimate | $\chi^2$ | Df | Estimate | $\chi^2$ | Df | Estimate | $\chi^2$ |
| ENVd | 1 | 0.15 | 1.26 | 1 | 9.65 | 0.7337 | 1 | 0.14 | 2.1102 | 1 | 0.05 | 2.88 |
| cluster | 2 | -1.66 | 16.75*** | - | - | - | - | - | - | - | - | - |
| site | 2 | -0.31 | 909.67*** | 2 | 1.90 | 1060.28*** | 2 | 1.90 | 610.87*** | 2 | -0.68 | 448.11*** |
| ENVd*cluster | 2 | -0.11 | 3.07 | - | - | - | - | - | - | - | - | - |
| ENVd*site | 2 | -0.16 | 1.97 | 2 | -0.15 | 2.5663 | 2 | -0.15 | 2.8086 | 2 | -0.05 | 2.62 |
| cluster*site | 4 | 2.97 | 56.51*** | - | - | - | - | - | - | - | - | - |

Significant values are indicated by \*\*\* ( $P$ -value <0.0001); ‡A Poisson error distribution was applied for count data.

Finlay-Wilkinson regression stability analyses, measuring the plasticity of *C. bursa-pastoris* accessions across common gardens, revealed that the EUR and ME accessions (mean *bi*=-2.2, SD=2.7 and, *bi*=2.2, SD=1.3, respectively) were significantly more responsive (i.e. plastic) to environmental changes than the ASI accessions (mean *bi*=0.45, SD=0.8) (all *P-values* were significant, Wilcoxon signed rank test, Figure 3). We further investigated the stability within each genetic cluster (Figure 3). FWR revealed a lack of responsiveness of the ASI accessions to environmental conditions regardless of the site; the ASI accessions were indeed stable across environments. In contrast, the EUR and ME genetic groups showed a high responsiveness to environmental conditions (Figure 3b). The European accessions performed well at both the Uppsala and Toronto common gardens but fitness decreased steeply in the Guangzhou common garden (Figure 3c). In contrast, the Middle Eastern accessions performed better at the Guangzhou common garden than at the Uppsala and Toronto common gardens (Figure 3d). These analyses agreed with the results presented above showing stable low fitness of ASI accessions across environments, high fitness of the EUR accessions in Northern Europe (Uppsala), and high fitness of the ME accessions in Southern China (Guangzhou). The FWR within the EUR genetic cluster also revealed that two accessions from Russia had negative regression slopes (Figure 4c) compared to the rest of the EUR cluster suggesting that although belonging to the same genetic group than European accessions, Russian accessions from Siberia might have started to diverge at the phenotypic level from their western counterparts (model [5]).

**Figure 3.**
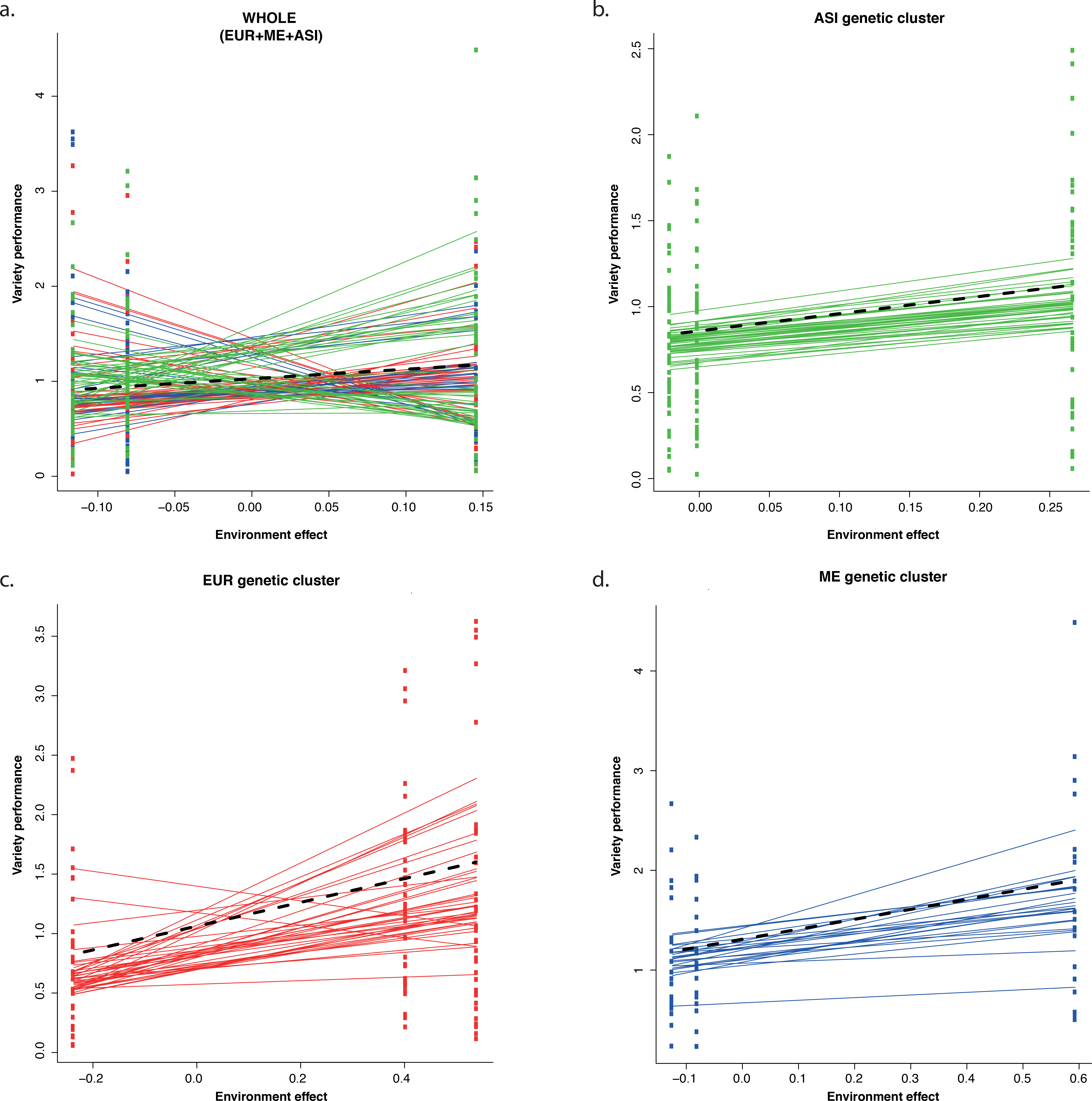
Finlay-Wilkinson Regression of accessions of *Capsella bursa-pastoris*. The y-axis is the fitness of individual accessions measured in each common garden and the x-axis is the environmental effect that is estimated by the mean of all accessions considered in the graphic in Uppsala, Guangzhou, and Toronto common gardens. The accessions included are from (a) the whole dataset (WHOLE) and accessions from the (b) Asian (ASI), (c) European (EUR) and (d) Middle Eastern (ME) genetic clusters. Each color represents a different accession. Lines are fitted values and circles are the cell means of genotype by environment combinations. The black dotted line is the average slope over all accessions.

**Figure 4.**
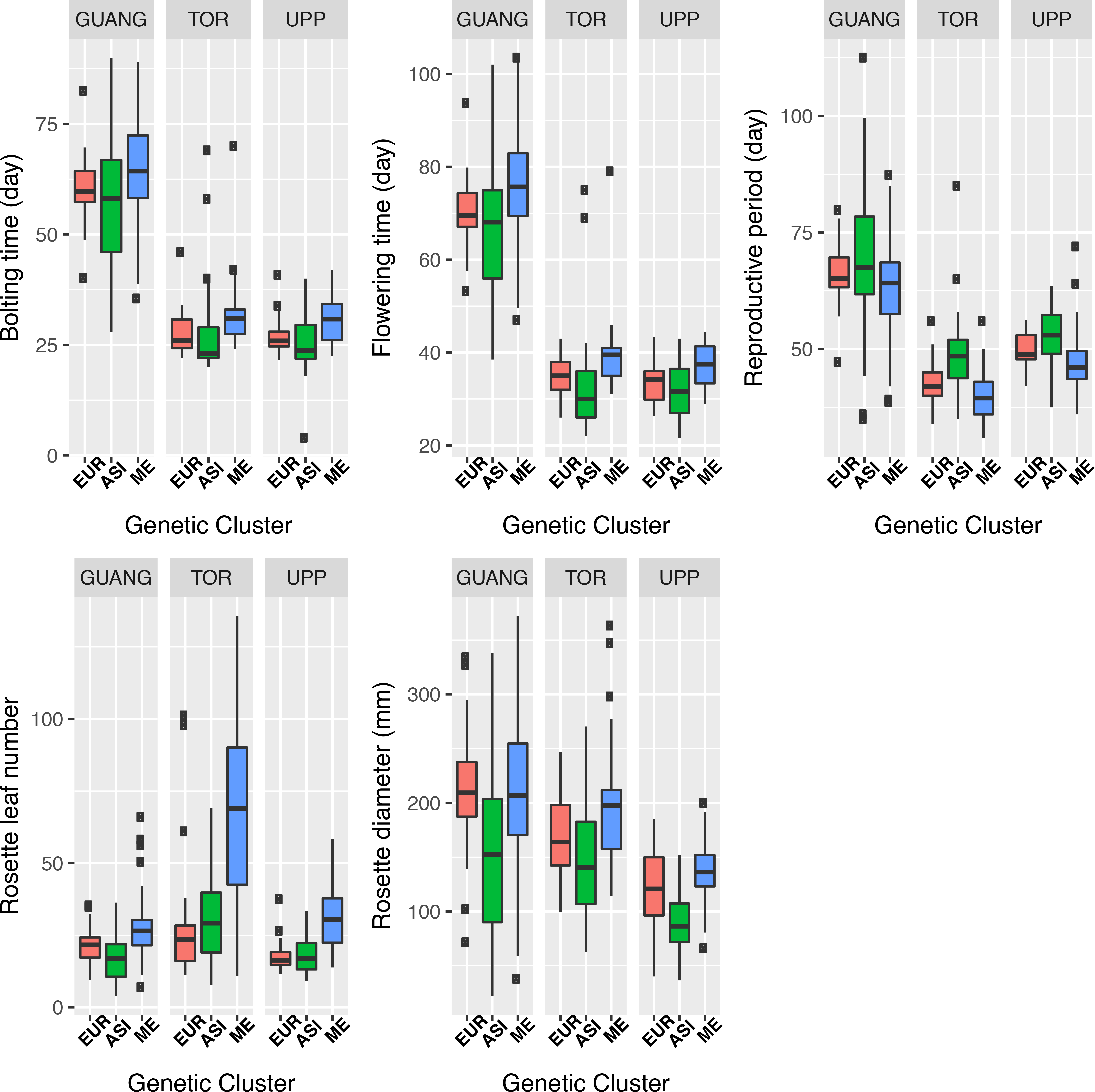
Boxplots of natural variation within each common garden (GUANG, TOR, UPP) for each genetic group (EUR, ASI and ME). BT: Bolting time, FT: flowering time, RP: reproductive period, MAXROS: Maximum rosette diameter, NUMBLEAVES: number of leaves of the rosette. Genetic cluster interactions were significant for the five phenotypic traits.

### 3. Genetic diversity and genetic distance from the common garden, and fitness

Fitness interpolations compared with genetic distance interpolations at the Guangzhou site (Figure 1B) suggested that the further genetically an accession is from the Guangzhou site, the better it fits the local environment, predominantly due to the ME accessions’ high performance in Guangzhou. This observation was statistically supported at the level of the entire dataset and within the EUR genetic cluster (model [3], *P* <0.001, Table 3). In contrast, at the Uppsala site, the correlation between genetic distance from the common garden and fitness of the accession was significantly negative within the ME genetic cluster.

**Table 3.** Effects of the genetic distance to the site (*GENDist*) and the environmental distance (*ENVd*) on the fitness of each individual (W) of *Capsella bursa-pastoris* at each common garden (Model [3]).

| Common garden | Explicative Variable | W_WHOLE‡ |  |  | W_ME‡ |  |  | W_EUR‡ |  |  | W_ASI‡ |  |  |
| --- | --- | --- | --- | --- | --- | --- | --- | --- | --- | --- | --- | --- | --- |
| | | Df | Estimate | $\chi^2$ | Df | Estimate | $\chi^2$ | Df | Estimate | $\chi^2$ | Df | Estimate | $\chi^2$ |
| Uppsala | ENVd | 1 | -0.04 | 0.32 | 1 | -0.03 | 0.07 | 1 | -0.12 | 0.76 | 1 | -0.13 | **4.06\*** |
|  | GENDist | 1 | 0.00 | 0.01 | 1 | -0.08 | **10.06\*\*\*** | 1 | 0.00 | 0.06 | 1 | 0.00 | 0 |
| Guangzhou | ENVd | 1 | 0.09 | 0.42 | 1 | 0.05 | 0.21 | 1 | -0.10 | 0.91 | 1 | 0.11 | 0.21 |
|  | GENDist | 1 | 0.02 | **5.57\*\*** | 1 | 0.02 | 0.87 | 1 | 0.04 | **7.07\*\*\*** | 1 | 1.58 | 0.87 |

Values are $\chi^2$ -values and significance from likelihood ratio tests of mixed-effect models; Significant values are marked with \*\*\* (*P-value* <0.001), \*\* (*P-value* <0.01) or \* (*P-value* <0.05); Df: degree of freedom; ‡A Poisson error distribution was applied for count data.

Within each common garden, the correlation between genetic diversity and the average fitness per population was positive but non significant, except for the Uppsala common garden which showed a strong ENVd*π effect (*P*=0.01) (model [4]).

### 4. Variation and adaptive values of phenotypic traits

The variation in phenology and growth among common gardens was significant (Table 4 and Figure 4), with higher variation in phenological than growth traits (Figure 4). Compared with the common gardens at Uppsala and Toronto, phenology (i.e. bolting time, flowering time and reproductive period) was delayed for all accessions at the Guangzhou common garden (Figure 4). The genetic cluster*common garden site interaction on phenotypic trait variation was also significant (Table 4), and plants from different genetic groups showed the significant differential phenotypic response to the different site characteristics (Table 4: Models [5] and [5’]).

**Table 4.** Effects of the common garden site (site), the genetic cluster (cluster: EUR, ME or ASI) on each phenotypic trait measured on the individuals of *Capsella bursa-pastoris* (Models [5] and [5’]).

|  | Model [5] |  |  |  |  |  | Model [5']: per genetic cluster |  |  |  |  |  |
| --- | --- | --- | --- | --- | --- | --- | --- | --- | --- | --- | --- | --- |
|  | ENTIRE |  |  |  |  |  | ASI |  | ME |  | EUR |  |
|  | Cluster (Df=2) |  | Site (Df=2) |  | Cluster*Site (Df=4) |  | site (Df=2) |  | site (Df=2) |  | site (Df=2) |  |
| | Estimate | $\chi^2$ | Estimate | $\chi^2$ | Estimate | $\chi^2$ | Estimate | $\chi^2$ | Estimate | $\chi^2$ | Estimate | $\chi^2$ |
| BT‡ | -0.02 | 17.9 | -0.77 | 17445.2 | 0.003 | 21.8 | -0.78 | 7579.2 | -0.77 | 3311.4 | -0.74 | 6707.6 |
| FT‡ | -3.5E-05 | 20.1 | -0.68 | 14740.7 | -0.02 | 16.3 | -0.72 | 7173.8 | -0.69 | 3039.8 | -0.69 | 4896.1 |
| RP‡ | 0.01 | 25.3 | -0.36 | 272.6 | 0.003 | 23.8 | -0.33 | 221.5 | -0.36 | 310.7 | -0.39 | 244.1 |
| ROSDIA | -1.47 | 38.3 | -2.68 | 384.4 | 1.25 | 54.8 | -1.08 | 350.5 | -2.68 | 258.1 | -1.40 | 130.3 |
| LEAVESNB‡ | -0.07 | 63.9 | -0.03 | 371.1 | 0.47 | 191.2 | 0.35 | 294.7 | -0.03 | 31.47 | 0.52 | 556.1 |

All *P-values* were highly significant. ‡A Poisson error distribution was applied for count data; BT: bolting time; FT: Flowering time; RP: reproductive period; LEAVESNB: rosette leaf number; ROSDIA: maximum diameter of the rosette; ASI: Chinese genetic cluster; EUR - European genetic cluster; ME - Middle Eastern genetic cluster; ENTIRE: Entire dataset; Df: degree of freedom; \* indicates interaction.

We explored the within-site selection differential. At the Guangzhou site, there was a significant genetic cluster effect in selection differential (*P*<0.0001, *χ^2^_179_*= 2.12). Actually, only the EUR genetic cluster showed significant positive directional selection for flowering time and bolting time, negative directional selection for reproductive period, and disruptive selection for the number of leaves. Therefore, at the Guangzhou site, the EUR accessions that bolted and flowered earlier, and had a longer reproductive period and higher fitness. At the Uppsala site, there was also a significant genetic cluster effect (*P*<0.0001, *χ^2^_210_*=1.17) indicating contrasted selection gradients between the ASI and the EUR accessions. The EUR accessions showed signatures of positive directional selection for bolting time and flowering time, stabilizing selection for reproductive period, and disruptive selection for rosette diameter. For the ASI genetic cluster, we found positive directional selection for flowering time, rosette diameter and number of leaves, and negative directional selection for reproductive period, suggesting that an ASI accessions with late flowering time, large rosette diameter and number of leaves, but a short reproductive period had higher fitness. At the Toronto site, we also found a significant genetic cluster effect (*P*=0.03, *χ^2^_91_*=0.54), with only the ME genetic cluster showing positive directional selection for rosette diameter and number of leaves. Therefore, the larger rosette diameter and number of leaves for ME accessions, the higher their fitness at the Toronto site (Table 5).

**Table 5.** Selection differential (*S*) for phenological and morphological traits for each common garden (Uppsala, Guangzhou, Toronto) in *Capsella bursa-pastoris*.

| Common garden | Trait | ENTIRE | ME | EUR | ASI |
| --- | --- | --- | --- | --- | --- |
| Guangzhou | BT | -0.23(0.16) | -0.09 (0.63) | **-0.72 (0.28)** | -0.04(0.17) |
|  | FT | -0.31 (0.18) | -0.28 (0.72) | **-0.85 (0.31)** | -0.03 (0.21) |
|  | RP | 0.19 (0.18) | 0.45 (0.79) | **0.78 (0.34)** | -0.01 (0.20) |
|  | ROSDIA | 0.09 (0.08) | 0.10 (0.23) | -0.01 (0.14) | 0.06 (0.09) |
|  | LEAVESNB | -0.09(0.08) | -0.13 (0.26) | -0.19 (0.12) | 0.03 (0.11) |
| Uppsala | BT | -0.15 (0.11) | 0.05 (0.38) | **-1.33 (0.24)** | 0.05 (0.13) |
|  | FT | **0.34 (0.14)** | 0.79 (0.41) | **-1.39 (0.37)** | **0.41(0.15)** |
|  | RP | **-0.57 (0.20)** | -1.06 (0.71) | 0.70 (0.45) | **-0.65 (0.23)** |
|  | ROSDIA | **0.47 (0.07)** | 0.27 (0.17) | 0.36 (0.23) | **0.54 (0.10)** |
|  | LEAVESNB | **0.21 (0.06)** | 0.22 (0.25) | -0.16 (0.13) | **0.29 (0.09)** |
| Toronto | BT | -0.01 (0.01) | -0.21 (0.02) | -0.27 (0.01) | -0.46 (0.01) |
|  | FT | 0.17 (0.13) | 0.79 (0.46) | -0.28 (0.24) | 0.068 (0.29) |
|  | RP | -0.31 (0.21) | -0.95 (0.58) | 0.44 (0.46) | 0.023 (0.34) |
|  | ROSDIA | **0.34 (0.09)** | **0.68 (0.31)** | 0.28 (0.16) | 0.22 (0.14) |
|  | LEAVESNB | **0.14 (0.05)** | **0.46 (0.20)** | 0.07 (0.08) | 0.12 (0.16) |

BT: bolting time; FT: Flowering time; RP: reproductive period; LEAVESNB: rosette leaf number; ROSDIA: maximum diameter of the rosette; ENTIRE: Entire dataset; ASI: Chinese genetic cluster; EUR - European genetic cluster; ME - Middle Eastern genetic cluster. Significant coefficients (*P-value* <0.001) are bolded; Values between brackets are the Standard Errors (SE).

**Table 6.** Gene ontology (GO) enrichment and flowering time candidate genes detected through SNP-environment associations in *Capsella bursa-pastoris*.

a) GO terms with significant enrichment in Asia and Europe
| Term | Count | % | *P*-value |
| --- | --- | --- | --- |
| **Asia** |  |  |  |
| GO:0008270~zinc ion binding | 17 | 19.32 | 3.64E-04 |
| GO:0000815~ESCRT III complex | 3 | 3.41 | 4.57E-04 |
| GO:0010008~endosome membrane | 3 | 3.41 | 0.003 |
| GO:0044440~endosomal part | 3 | 3.41 | 0.003 |
| **Europe** |  |  |  |
| GO:0016787~hydrolase activity | 27 | 25.23 | 0.013 |
| GO:0042221~response to chemical stimulus | 14 | 13.08 | 0.044 |
| GO:0010038~response to metal ion | 5 | 4.68 | 0.083 |
| GO:0010038~response to metal ion | 5 | 4.68 | 0.090 |

b) Flowering time candidate genes detected through SNP-environment associations, with abbreviated TAIR descriptions
| Gene | Function | Region |
| --- | --- | --- |
| AT1G17760<br>ATCSTF77 | Mediates<br>silencing of FLC | Asia |
| AT5G65070<br>AGL69 | FLC paralog | Asia |
| AT1G63030<br>DDF2 | Transcription<br>factor involved<br>in gibberellin<br>regulation | Europe |
| AT4G35900<br>FD | Positive<br>flowering time<br>regulator | Europe |

Regression analyses allowed the detection of a significant selection gradient for each trait in at least one of the three sites and genetic clusters. However, we detected selection for different traits depending on the dataset used (i.e. entire, EUR, ME, ASI, Tables 5 and S3). The total variance in fitness among accessions using the entire dataset (257 individuals in total) was best explained by phenological traits in the Uppsala site: *R*^2^_(*Uppsala*)_=0.28, *R*^2^_(*Guangzhou*)_=0.24, *R*^2^_(*Toronto*)_=0.10. However, the total variance was globally low for each common garden. In Uppsala, we found positive directional selection for flowering time, rosette diameter and number of leaves, and negative directional selection for reproductive period. This suggests that accessions at the Uppsala site, that flowered later and had larger rosettes and number of leaves, and had shorter reproductive period were better fitted to the Northern European environmental conditions. At the Toronto site, we found positive directional selection for rosette diameter and number of leaves, suggesting that accessions that had larger rosettes and higher number of leaves had higher fitness.

Overall, these results suggested a latitudinal selective gradient effect: accessions that flowered, bolted earlier, and with a longer reproductive period had a higher fitness in Southern China (i.e. Guangzhou), while the fitness of accessions with delayed bolting and flowering times and a shorter reproductive period was higher in Northern Europe (i.e. Uppsala). At the Toronto site, we also showed that the significant selective gradient was mainly driven by the ME genetic cluster.

We carried out multiple linear regressions to estimate linear regression coefficient (*β*) and quadratic selection coefficient (γ) (Table S3). Within the entire dataset, there were contrasted patterns of selective gradients among the three common gardens. There was evidence for stabilizing and disruptive selection for rosette diameter and number of leaves respectively at the Guangzhou common garden. This may seem surprising, but the stabilizing or disruptive nature of selection also depends on where the phenotypes are on the fitness curve. One can have a quadratic fitness function but only be in a monotonic part of this function if the observed range of phenotypes does not include the minimum or maximum of the quadratic function. It is maybe the case here.

### 5. Association of SNPs with environmental variables

Environmental associations with SNP frequencies also reflected limited adaptation throughout the sampled range with region of origin shaping its genetic basis. Consistent with weak adaptive differentiation, we did not observe significant isolation by environment in any sample (Mantel test, p>0.05). However, through LFMM we identified 142 and 105 SNPs with significant associations in Europe and Asia, respectively, using the top 1% most significant associations. These SNPs overlapped substantially with the top 5% of SNPs identified when we repeated this analysis with only one sample from each population (68.9% overlap in Europe, 80.5% overlap in Asia). We detected two flowering time candidate genes in each of Europe and Asia (Table 6). In Asia, we found two genes involved in the FRI and FLC vernalization pathway (i.e. ATCSTF77 and AGL69, Table 6), while in Europe, we found the gibberellin acid response factors DDF2 and the daylight independent flowering time regulator FD.

The presence of flowering time candidate genes aligned well with our observation of directional selection in flowering time in both regions, and suggested subtle environmental adaptation may occur in *C. bursa-pastoris*.

According to the enrichment analysis, SNPs with potential environmental associations in Europe and Asia were distinct. Europe and Asia only shared two putatively adaptive LFMM SNPs (*P*=0.84). A total of 11 out of a potential 177 (13.5%) combined unique genes that contained SNPs with significant associations (overlap *P*=0.11). Environmentally associated SNPs showed significantly higher mean pairwise *r^2^* both within (*P*<0.001) and across chromosomes (*P*<0.001) compared to 1,000 random draws of SNPs from the corresponding dataset in Europe indicating strong LD among putatively adaptive SNPs. In contrast, LD among putatively adaptive SNPs in Asia was not significantly higher within or among chromosomes compared to 1,000 random samples (*P*=0.81, *P*=0.56). Estimated effect sizes for SNPs with significant associations were also significantly higher in Asia than Europe (Figure 5, Student’*s-t*=-3.55, *P*<0.0004). Furthermore, we also found an excess of alleles with MAF from 0.3-0.4 compared to the MAF distribution of sites with significant LFMM associations from 500 permutations of the environmental data matrix in Europe (*P*<0.0001) but not Asia, consistent with standing variation predominantly driving adaptation in Europe and playing a lesser role in Asia (Figure 6). Together, these results suggest adaptation may have a different genetic basis in Europe and Asia: many linked SNPs of small effect segregating from standing variation may contribute to adaptation in Europe, and while associations occur through unlinked larger effect SNPs that more often involve new mutations in Asia.

**Figure 5.**
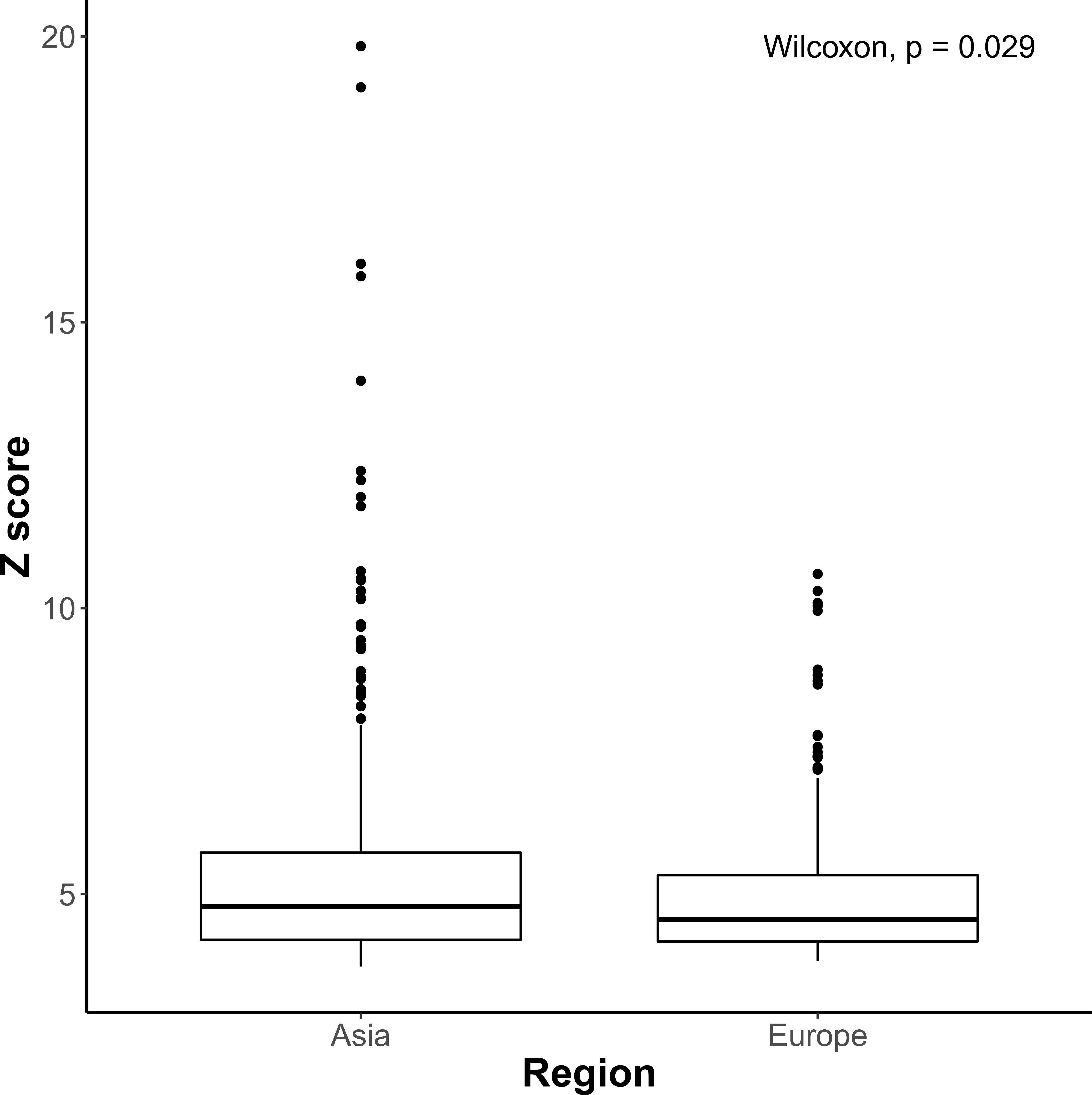
Boxplot of Z-scores of SNPs with significant associations in Asia and Europe from Latent Factor Mixed Model (LFMM) approach for *Capsella bursa-pastoris*. Z-scores are standardized *β* terms from the LFMM mixed model regression (Frichot *et al.* 2013).

**Figure 6.**
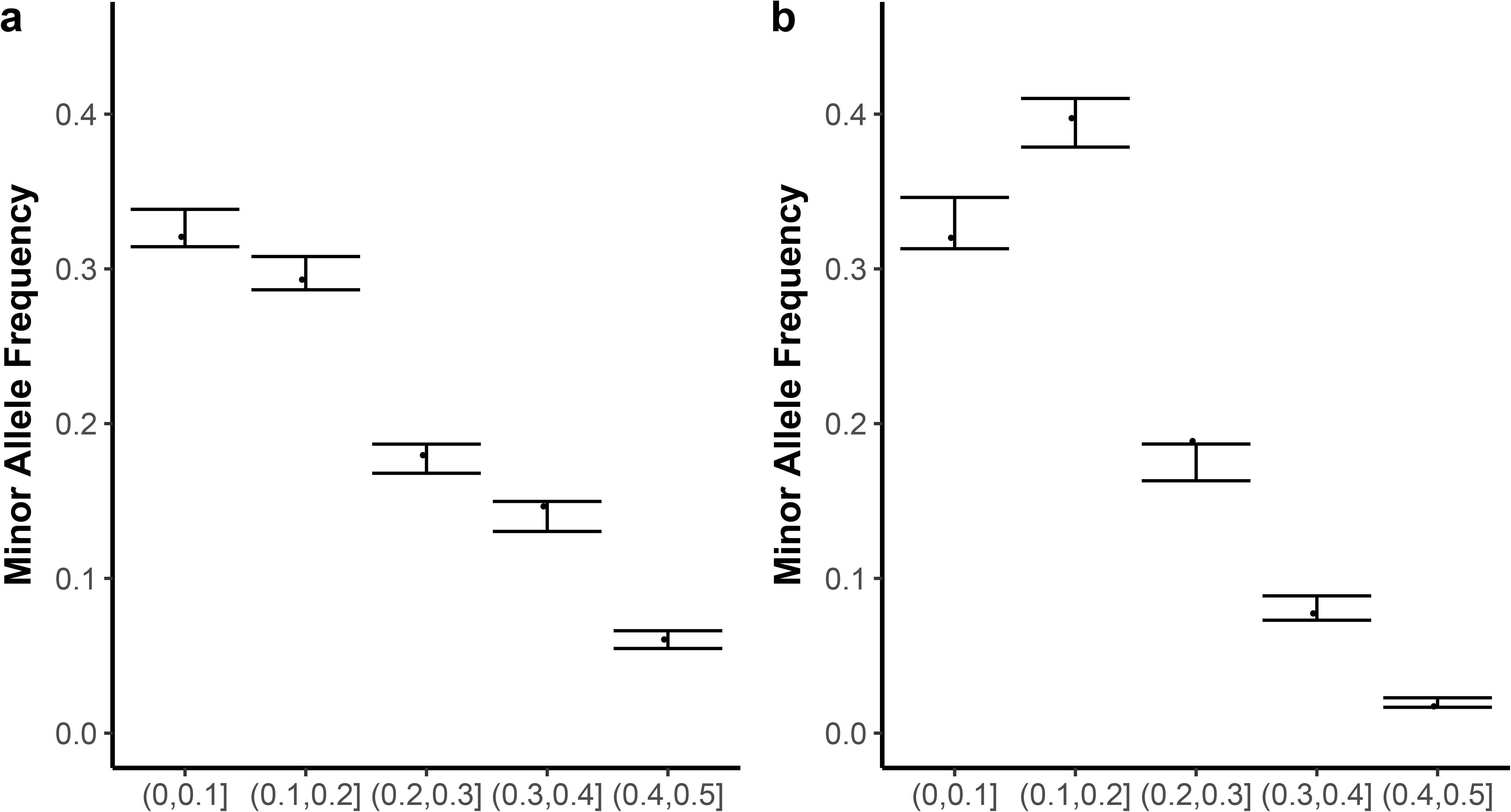
Minor allele frequency spectra of SNPs with significant environmental associations in *Capsella bursa-pastoris* in Asia (A) and Europe (B) with 95% confidence intervals based on 500 permutations.

## Discussion

In the present study, we installed common garden experiments on three continents to assess the fitness and levels of local adaptation of the shepherd’s purse, *C. bursa-pastoris*. Patterns of local adaptation were generally weak, and there were even indications of maladaptation in the large group of Asian accessions. The latter was characterized by low diversity at the genome level as well as for traits such as responsiveness to environmental changes. The selection gradients on some of the traits related to fitness were of opposite sign in Guangzhou, and in Uppsala and Toronto. On the other hand, the association between the molecular polymorphism and environmental variables that are suggestive of local adaptation were detected in both Europe and Asia.

These results have important consequences for our understanding of the evolution of selfers and, in particular, the evolution of polyploid selfers. Following early suggestions by George Shull (Shull 1929), who, by and large, initiated research in the *Capsella* genus, one of the main initial aims of the research program to which the current study belongs was to explain the success of *C. bursa-pastoris* compared to its diploid relatives. On the face of it, *C. bursa-pastoris* does look successful as it has one of the largest geographical ranges among plants and certainly much larger than the ranges of its three diploid relatives. But is this success long lasting or is it ephemeral? Before addressing this question we shall discuss some strengths and caveats of the current study.

### Possible caveats

The standard experimental design to measure local adaptation in plants is reciprocal transplant. It is a straightforward and powerful design but it is generally limited to a couple of sites. As such, it will fail to provide an estimate of local adaptation if the latter is viewed, as in the present study, as a property of a metapopulation (Blanquart *et al*. 2013). Common gardens are an alternative to reciprocal transfer experiments. Clearly common gardens provide a more indirect measure of local adaptation but they can accommodate a much larger number of samples and will therefore be better at capturing the metapopulation’s local adaptation. They might also better account for the species’ demographic history.

In the present experiment, we used three common garden experiments, two within the native range of *C. bursa-pastoris*, and one outside of it. To estimate local adaptation, we included an environmental distance effect between the sampling locations and the common garden locations. In our analysis, we then interpreted the effect of environmental distance as a measure of the strength of local adaptation and the interaction between environmental distance and genetic cluster as measuring differences of local adaptation among the three genetic clusters.

Clearly, the step of adding environmental distance into the model, compared to a standard reciprocal transfer experiment, complicates matters and could explain the apparent lack of local adaptation observed in the present study. However, we do not think this is the case. First, if there were a strong local pattern effect, we would have already detected it simply by plotting the extrapolated fitness onto a map. Second, the environmental distance was calculated based on 19 bioclimatic variables from the Worldclim database that should capture fairly well the factors that differentiate the experimental sites. Apart from climatic conditions, other conditions at each experimental site were quite similar and this could also have contributed to the lack of strong signature of local adaptation, although it is doubtful that other environmental conditions than climate contribute to local adaptation in a weed like the shepherd’s purse at such a large geographical scale. To the best of our knowledge, there has not been a general study of the local ecological niche of the shepherd’s purse. However, our own observations and the few studies available, in particular the study by Caullet (2011) (Caullet 2011) that exhaustively studied populations of *C. bursa-pastoris* and *C. rubella* across a 4 km^2^ agricultural landscape of Central France over two years, suggest that *C. bursa-pastoris* may be actually specific to highly disturbed environments (hedges of fields, for instance). However, within such environments it does not seem to be confined to specific types of soil (Aksoy *et al*. 1998; Neuffer *et al*. 2018). The large spectrum of climatic environments covered by the accessions included in the experiment also suggests that insufficient sampling of the accessions is unlikely to explain the lack of a strong local adaptation pattern.

### A lack of local adaptation?

The apparent lack of local adaptation observed in the present study at the phenotypic level can be related to the results obtained by Kryvokhyzha et al. (2016). Using RNA-Seq data from 24 accessions originating from the three clusters, Kryvokhyzha and co-authors showed that the genes that were significantly differentially expressed between the three clusters, were no longer so once one had corrected the data for the genetic relationship between the three clusters. The differentiation could then have been mostly neutral or both neutral and adaptive in which case demographic history and adaptive differentiation were highly correlated and correcting for the first one would remove the effect of the latter. The fitness results obtained here would support at least a significant contribution of neutral processes. As in the RNA-Seq experiment, the present data does not rule the presence of significant local adaptation at a more local scale. However, even at that scale local adaptation does not seem to be pronounced in *C. bursa-pastoris* (Caullet 2011).

It therefore seems likely that *C. bursa-pastoris* might have been able to cope with diverse environments primarily through phenotypic plasticity. Consistent with this hypothesis, populations with higher plasticity (from the EUR and ME clusters) also showed the highest fitness in gardens both within and outside of the native range. Caullet and coauthors (2011) also found plasticity best explained fitness across soil niches when they planted both *C. bursa-pastoris* and *C. rubella* in four different disturbed environments where both species are naturally found: along paths, hedges of the paths, or within a transition zone between paths and fields and inside the internal hedge of the fields. The transplant environment did not significantly affect the fitness of *C. bursa-pastoris*. As a whole, in this field experiment proposed by Caullet and coauthors phenotypic plasticity was higher in *C. bursa-pastoris* than in *C. rubella* but there was no evidence of strong local adaptation in *C. bursa-pastoris*. On the other hand, we cannot completely rule out the presence of local adaptation as significant associations between polymorphisms and environmental variables were observed across the whole range, in particular in Asian populations where otherwise signs of maladaptation were the most pronounced.

### Local adaptation in an expanding self-fertilizing species

Two major differences between Asian accessions and European and Middle Eastern ones are a higher rate of deleterious alleles, as well as a lower genetic diversity in the former than in the latter, and recent introgression of the former by *Capsella orientalis* (Kryvokhyzha *et al*. 2017). These differences in nucleotide polymorphism data were reflected by similar tendencies at the phenotypic level: the Asian accessions have the lowest fitness in two environments and a much lower fitness than the Middle Eastern accessions in the Chinese site itself. Their response to the environment was also markedly different from the other two populations with less variation among accessions and a lower response to environmental change. We also previously showed in a competition experiment where different accessions were subjected to different competition pressures that Chinese accessions were more sensitive to competition than European or Middle Eastern ones (Yang *et al*. 2018). All these characteristics are consistent with a recent bottleneck in the history of the Chinese population leading to a lower fitness. Does this imply a lack of ability to respond to selection and adapt locally? Probably not, as we did observe evidence of local adaptation at the molecular level as well as significant selection gradients for rosette size and flowering time in ASI. One possible explanation to this apparent contradiction is that Chinese populations are far from demographic equilibrium and far from their fitness optimum. So even though they carry more deleterious mutations than their European or Middle East relatives and have overall a lower fitness, positive mutations of large effect are also more likely to occur and go to fixation than in populations close to their optimum where such mutation will be rare (Fisher 1930). Similarly, in humans, while Europeans and Asians also went through a bottleneck, it did not seem to have impaired drastically their ability to respond to selection compared to the more diverse African populations, even if selection is weak and neutral processes important in determining the fate of selected alleles (Coop *et al*. 2009; Granka *et al*. 2012). Genomic associations may then reflect subtle, ongoing adaptation to factors outside of the environmental conditions measured in this study.

Finally, Kryvokyhyzha et al. (2017) detected introgression of *C. orientalis* into Chinese accessions of *C. bursa-pastoris. Capsella orientalis* has particularly low genetic diversity and a high number of deleterious alleles. Hence the recent expansion of *C. bursa-pastoris* and the recent gene exchanges from a parental species could also have contributed to the pattern of maladaptation observed here. Under this scenario, the Chinese accessions inherited and still have not purged the deleterious alleles introgressed from its parental species.

### Phenology and growth traits relate to relative fitness of plants at the different latitudes

We characterized the nature of phenotypic variation and adaptation among the three genetic clusters along a latitudinal environmental gradient and the nature of selection on traits. We found evidence that both phenology and growth traits relate to relative fitness of plants at the different latitudes. In Guangzhou, accessions that flowered early had higher fitness while the converse was true in Uppsala. However, this general pattern varied across clusters and was not consistent. Overall, it therefore seems that the variation in fitness is not explained by past natural selection and most of it could simply reflect phenotypic plasticity. It should be noted, however, that the conditions under which the plants were grown in the present experiments do not reflect those that prevail under natural conditions where, for instance, gardening and farming activities may favor early flowering genotypes (Neuffer & Hurka 1986). A different clustering of the populations and different environmental conditions might have led to the emergence of a clearer pattern of selection on fitness components than a clustering based on geography coupled with uniform and favorable common garden conditions.

## Conclusion

The present multisite common garden experiment confirms previous studies suggesting that demography, in particular, founder events and introgression, played a major role in shaping the current genotypic and phenotypic diversity in *C. bursa-pastoris* (Neuffer & Albers n.d.; Kryvokhyzha *et al*. 2016, 2017). We also showed that local adaptation, if it occurs in *C. bursa-pastoris*, occurs at a different geographical scale and under different environmental conditions than the ones used here.

## Acknowledgments

We thank the Swedish Research Council and the Erik Philip Sörensen Foundation (to ML), Uppsala University and EMBO Short-Term Fellowships fund research exchanges (to AC) for funding. This research was also funded by the General Program of National Natural Science Foundation of China (grant No. 31370005, to HRH). AS was supported by an NSERC CGS-M and an Ontario Graduate Scholarship. We thank Julia Dankanich, Josefine Stångberg, Johanna Nyström, Sara Kurland and Uriel Gélin for field assistance in Uppsala; Lin-Lin Wu, Zhi-Bing Xie, Qiu-Ling Guan, Gui-Yu Lin for field assistance in Guangzhou; and Viktor Mollov and Niroshini Epitawalage for field assistance in Toronto.

